# Dependence of Intravoxel Incoherent Motion diffusion MR threshold *b*-value selection for separating perfusion and diffusion compartments and liver fibrosis diagnostic performance

**DOI:** 10.1101/164129

**Authors:** Yao Li, Pu-Xuan Lu, Hua Huang, Jason Leung, Weitian Chen, Yi-Xiang Wang

## Abstract

**Purpose:** To explore how the selection of threshold *b*-value impacts Intravoxel Incoherent Motion (IVIM) diffusion parameters of PF (*f*), D_slow_ (*D*),and D_fast_ (*D\**) values and their performance for liver fibrosis detection.

**Materials and Methods:** Fifteen healthy volunteers and 33 hepatitis-b patients were included. With a 1.5 T MR scanner and respiration gating, IVIM data was acquired with 10 *b*-values of 10,20,40,60,80,100,150, 200, 400, and 800 s/mm^2^. Signal measurement was performed on right liver. Segmented-unconstrained analysis was used to compute IVIM parameters, and six threshold *b*-values between 40 and 200 s/mm^2^ were compared. PF, Dslow, and Dfast values were placed along the *x*-axis, *y*-axis, and *z*-axis, and a plane was defined to separate volunteers from patients.

**Results:** Higher threshold *b*-values were associated with higher PF measurement; while lower threshold *b*-values led to higher Dslow and Dfast measurements. The dependence of PF, Dslow, and Dfast on threshold *b*-value differed between healthy livers and fibrotic livers; with the healthy livers showing a higher dependence. Threshold *b*-value=60 s/mm^2^ showed the largest mean distance between healthy liver datapoints vs. fibrotic liver datapoints in 3-dimensional space.

**Conclusion:** For segmented-unconstrained analysis, the selection of threshold *b*-value=60 s/mm^2^ improves IVIM diffusion differentiation between healthy livers and fibrotic livers.

## Introduction

Viral hepatitis is the most common blood-borne infection worldwide [1,2]. Chronic viral hepatitis can lead to hepatic fibrosis, cirrhosis and hepatocellular carcinoma [3]. Clinically liver fibrosis usually has an insidious onset and progresses slowly over decades. A noninvasive and quantitative technique for detecting liver fibrosis is highly desirable. Treatment with combined therapies on underline etiology and fibrosis simultaneously can expedite the regression of liver fibrosis and promote liver regeneration [4]. Le Bihan *et al* [5,6] proposed the principle of intravoxel incoherent motion (IVIM) diffusion weighted MRI (DWI) which enables the quantitative parameters that separately reflect tissue diffusivity and tissue microcapillary perfusion to be estimated. IVIM signal attenuation is modeled according to the equation

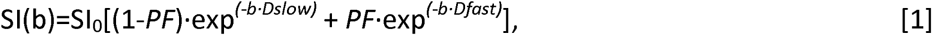

where SI(b) and SI_0_ denote the signal intensity acquired with the *b*-factor value of *b* and *b*=0 s/mm^2^, respectively. Perfusion fraction (PF, or *f*) is the fraction of the pseudo-diffusion linked to microcirculation, Dslow (or *D*) is the true diffusion coefficient representing the pure molecular diffusion (slow component of diffusion), and Dfast *(*D\**)* is the pseudo-diffusion coefficient representing the incoherent microcirculation within the voxel (perfusion-related diffusion, or fast component of diffusion) [7].

Since the initial liver IVIM DWI study of Yamada *et al* [8], there has been great interest of using this technique to study liver diseases such as liver fibrosis. Molecular water diffusion in fibrotic liver would be restricted by the presence of collagen fibers in the distorted lobular structure. Liver fibrosis is also known to be associated with reduced liver perfusion [9, 10]. However, IVIM parameters suffer from unsatisfactory scan-rescan reproducibility, and most reports demonstrated IVIM approach was unable to detect early liver fibrosis reliably [11]. More recently, with a dataset composed of 16 healthy subjects and 33 liver fibrosis patients ( *.XXX. 2012/2013 ivim dataset),* Wáng *et al* reported that a combination of Dslow, PF and Dfast, could be used to separate even F1 fibrotic livers from healthy livers [12]. The findings of this study are important as till now there is no reliable noninvasive method, being imaging or serum biomarkers, can reliably detect early stage liver fibrosis [13].

In Wáng *et al*’s analysis [12], the threshold *b*-value chosen was 200 s/mm^2^ and the segmented-unconstrained approach was used [14]. The segmented-unconstrained analysis remains the most commonly used method in literature [11, 14, 15]. Recently Park *et al* [15] described that segmented-fitting should be preferred in the cases of a limited number of *b*-values and a limited signal-to-noise ratio. *b*-value=200 s/mm^2^ has been commonly selected as a threshold value as perfusion component's influence on signal decay can be neglected for *b*-values ≥200 s/mm^2^ [14 -22]. However, with analysis of data sampled from healthy subjects, Wurnig *et al* [23] proposed threshold *b*-value of 20 s/mm^2^ and 40 s/mm^2^ for liver right and left liver robe respectively. Other threshold *b*-values have also been used in literature [24, 25]. Tissue parameters derived from IVIM analysis depend on the threshold *b*-value [23]. The optimal threshold *b*-value for liver IVIM analysis is yet undecided. Particularly, till now there is no empirical validation with data containing patients. In the current study, we explored how the selection of threshold *b*-value impacts PF, Dslow, and Dfast values; and more importantly, how threshold *b*-value impacts IVIM technique’s performance for liver fibrosis detection. As the IVIM diffusion weighted imaging (DWI) sequence is widely available in clinical MR scanners, it represents a promising alternative to existing techniques for liver fibrosis evaluation.

## Material and Methods

The *Shenzhen 2012/2013 ivim dataset* was acquired during the period from Aug 1, 2012 to Aug 15, 2013 [12, 19]. The study was approved by the institutional ethical committee, and the informed consent was obtained. With the initial dataset were screened for image quality, the IVIM images of two volunteer and one patient were adjudged to contain substantial motion artifacts and were therefore excluded for analysis (see supplementary document S1). Fifteen healthy volunteers (10 males, 5 females, mean age: 35.3-yrs old; range: 21–77 yrs old) and 33 consecutively viral hepatitis-b patients were included in the current study. The patient cohort had 15 stage F1 subjects (mean age: 31.8 yrs, 22-53 yrs) and 18 stage F2-4 subjects (mean age: 42 yrs, range: 22-53 yrs). The histology diagnosis for liver fibrosis followed the METAVIR score [26]. F0 and F1 livers are commonly referred to as without significant hepatic fibrosis; hepatic fibrosis (F2, F3, and F4) are commonly referred to as significant hepatic fibrosis, and F4 is also referred as cirrhosis [27]. Hepatic fibrosis is considered clinically significant if defined as F2 or greater than F2, and requiring medical attention [27, 28].

MR imaging was performed with a 1.5-T magnet (Achieva, Philips Healthcare, Best, Netherlands). In addition to standard anatomical liver imaging, the IVIM DWI sequence was based on a single-shot DW spin-echo type echo-planar imaging sequence, with ten *b*-values of 10, 20, 40, 60, 80, 100, 150, 200, 400, 800 s/mm2. SPIR technique (Spectral Pre-saturation with Inversion-Recovery) was used for fat suppression. Respiratory-gating was applied in all scan participants and resulted in an average TR of 1500 ms, and the TE was 63 ms. Other parameters included slice thickness =7 mm, matrix= 124×97, FOV =375 mm×302 mm, NEX=2, number of slices =6.

As the left lobe of liver is more likely to suffer from artifacts associated with the cardiac motion and B_0_ inhomogeneity susceptibility due its proximity to the stomach, in the current study only the right lobe of liver was measured. The ROIs in this study were re-drawn, independent from Wáng *et al*'s analysis, by a trained biomedical engineering graduate student (Y.T.L). The ROIs were positioned on *b*-value=10 DW images to cover a large portion of right liver parenchyma while avoiding large vessels and same ROI masks were propagated to all *b*-values images. All 6 slices per subject were evaluated, while the slices with notable motion artifacts and those demonstrates notable signal vs. *b*-value relationship outlier were then discarded. Finally, the slice used for final analysis varied between two to five slices (average: three slices).

The IVIM signal attenuation was modeled according to the Equation [1]. The estimation of Dslow was obtained by a least-squares linear fitting of the logarithmized image intensity at the threshold and higher *b*-values to a linear equation. The fitted line was then extrapolated to obtain an intercept at *b*-value=0. The ratio between this intercept and the SI_b=10_ gave an estimate of PF. Finally, the obtained Dslow and PF were substituted into Eq. [1] and were nonlinear least-square fitted against all *b*-values to estimate Dfast using the Levenberg-Marquardt algorithm.

Six threshold *b*-values, i.e. 40, 60, 80, 100, 150, and 200 s/mm^2^, were tested. For example, if threshold *b*-value was chosen to be 40 s/mm^2^, then *b*-values of 40 – 800 s/mm^2^ were used to compute Dslow. If threshold *b*-value was chosen to be 200 s/mm^2^, then *b*-values of 200, 400, and 800 s/mm^2^ were used to compute Dslow. For ROI analysis, the IVIM parameters were calculated based on the mean signal intensity of the whole ROI. All curve fitting algorithms were implemented in an accustom program developed on MatLab (Mathworks, Natick, MA, USA).

A three-dimensional tool was programed using IBM SPSS 23 for Windows (SPSS Inc., Chicago, IL, USA), and the measures of Dslow, PF, and Dfast were placed along the x-axis, y-axis, and z-axis. Data points from healthy volunteer were labeled as blue in the 3-dimensional space, F1 patients labeled as pink, and F2-4 patients labeled as red. Attempts were then made to separate healthy volunteers from all patients (F1-F4); healthy volunteers from significant patients (F2-F4). In addition, the Support Vector Machine (SVM) approach was used to quantitatively separate the F0 from F1-F4, or F0 from F2-F4 (29). SVM was used to find a plane (parametrized as Ax+By+Cz+D = 0) that was able to separate the data points into two groups. The distance of the closest data point from an individual group to the separating plane was defined as *d_i_*, where *i* represents the index of the group. The SVM algorithm was used to find an optimal plane which maximizes the margin defined as *d_1_* + *d_2_*. Prior to calculate the distance *d_i_*, the measured Dslow, PF, and Dfast are normalized by the following equation.

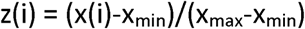

where x(i) is the original data and z(i) is the normalized data; x_max_ and x_min_ are the maximum and the minimum value of x(i), respectively. Note the range of z(i) after normalization is from 0 to 1 for each dimension.

## Results

For both healthy livers and fibrotic livers, higher threshold *b*-values were associated with overall higher PF measurement; while lower threshold *b*-values led to overall higher Dslow measurement and higher Dfast measurement (Fig 1, Fig 2). Liver fibrosis was progressively associated with a reduction of PF, Dslow, and Dfast (Fig 2). The dependence of PF, Dslow, and Dfast on threshold *b*-value differed between healthy livers and fibrotic livers; with the healthy livers showing a higher degree of dependence (Fig 2). With the *b*-value range between 40-200 s/mm^2^, PF value derived from threshold *b*-value=200 s/mm^2^ allowed the best separation of healthy liver vs fibrotic livers; while Dslow and Dfast derived from threshold *b*-value=40 s/mm^2^ allowed the best separation of healthy liver vs. fibrotic livers (Fig 2).

**Figure 1.**
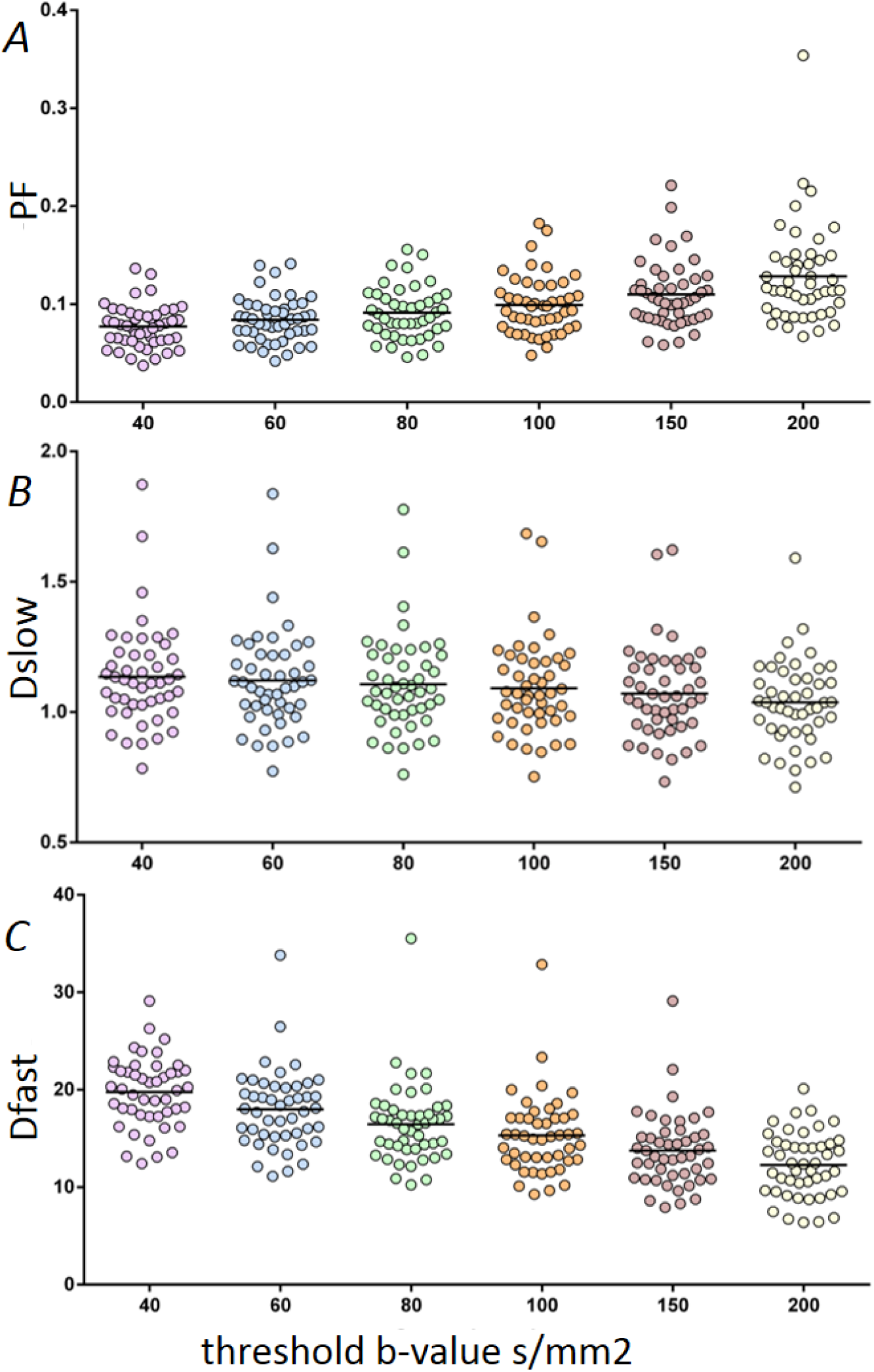
Values of PF, Dslow, and Dfast derived from different threshold b-values. All healthy livers (n=15) and fibrosis livers (n=33) were included. It can be seen that as the threshold b-value increases, PF value increases (A) while Dslow value (B) and Dfast value (C) decrease.

**Figure 2.**
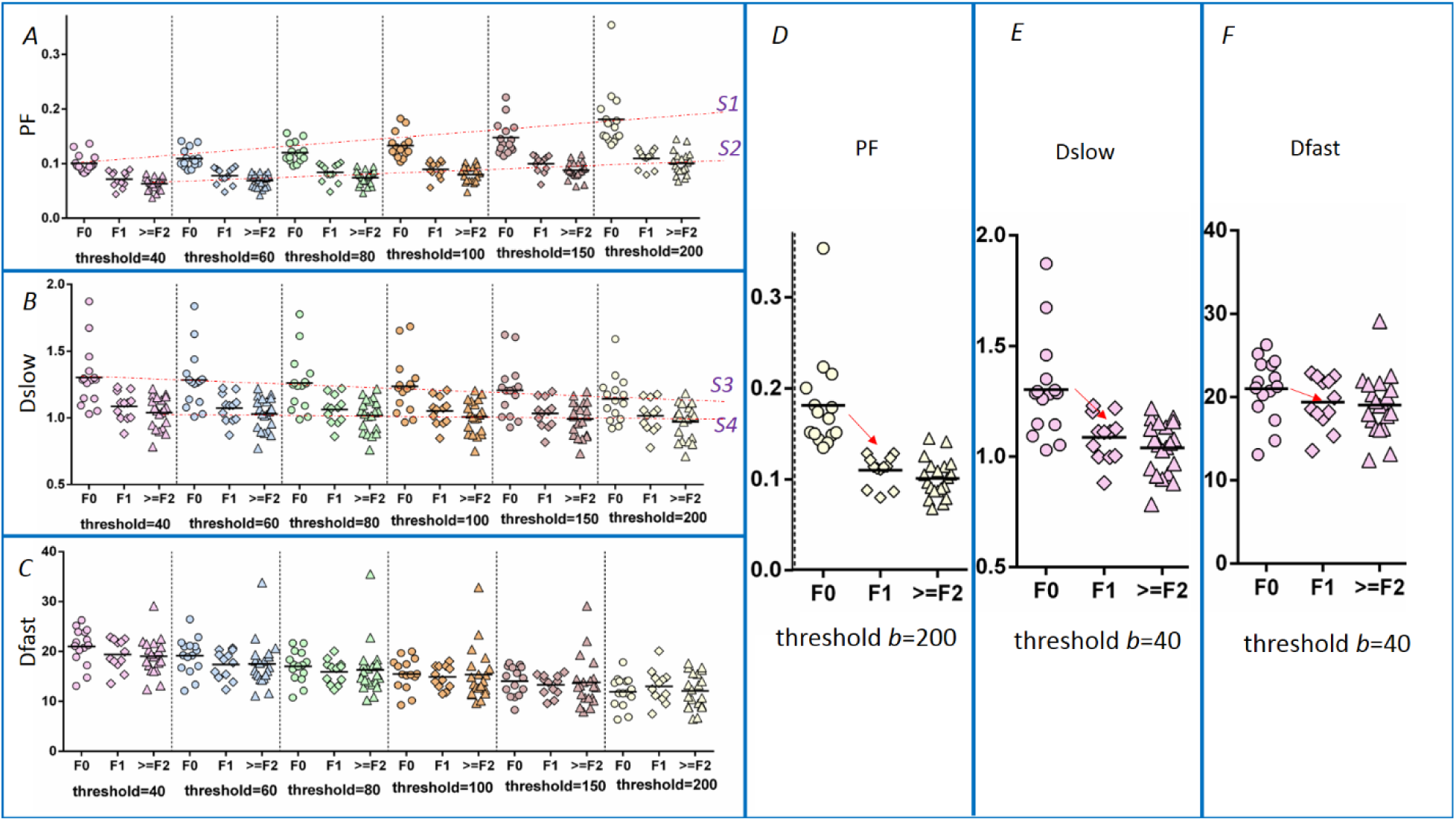
A comparison of healthy livers (F0), F1 stage livers, and F2-4 stage livers show liver fibrosis is progressively associated a reduction of PF (D), Dslow (E), and Dfast (F). The healthy livers show a higher threshold *b*-value dependence (e.g., slope of line S1> slope of line S2, slope of line S3> slope of line S4). PF value derived from threshold *b*-value=200 allows the best separation of healthy liver vs. fibrotic livers (A); while Dslow and Dfast derived from threshold *b*-value=40 s/mm^2^ allows the best separation of healthy liver vs. fibrotic livers (B, C). S1: estimated fit of dependence of PF on threshold *b*-value for healthy livers, S2: estimated fit of dependence of PF on threshold *b*-value for F2-4 livers, S3: estimated fit of dependence of Dslow on threshold *b*-value for healthy livers, S4: estimated fit of dependence of PF on threshold *b*-value for F2-4 livers, unit of PF: %, unit of Dfast and Dslow: mm^2^/s, unit of *b*-value: s/mm^2^.

The mean distance between healthy liver datapoints and fibrotic liver datapoints in 3-dimensional space is shown in figure 3. Compare to other threshold *b*-values, selection of *b*-value= 60 s/mm^2^ provided the largest distance, i.e. best separation between healthy livers vs.fibrotic livers. Compared with the previously used threshold *b*-value=200 s/mm^2^, *b*-value=60 s/mm^2^ increased the distance from 0.246 to 0.454 (84.5% increase, in RU (relative unit)). If the 1st˜10th pairs of patient and volunteer with closest distance were considered, the results remained the same to suggest that threshold *b*-value=60 s/mm^2^ provided the greatest distances (Fig 4). If only considering the separation of healthy livers (F0) vs. significant fibrotic livers (F2-4), then threshold *b*-value=80 s/mm^2^ provided results slightly better separation than *b*-value=60 s/mm^2^, and increased the distance from 0.359 to 0.594 (RU, 65.4% increase compared with *b*-value=200 s/mm^2^) (Fig 5).

**Figure 3.**
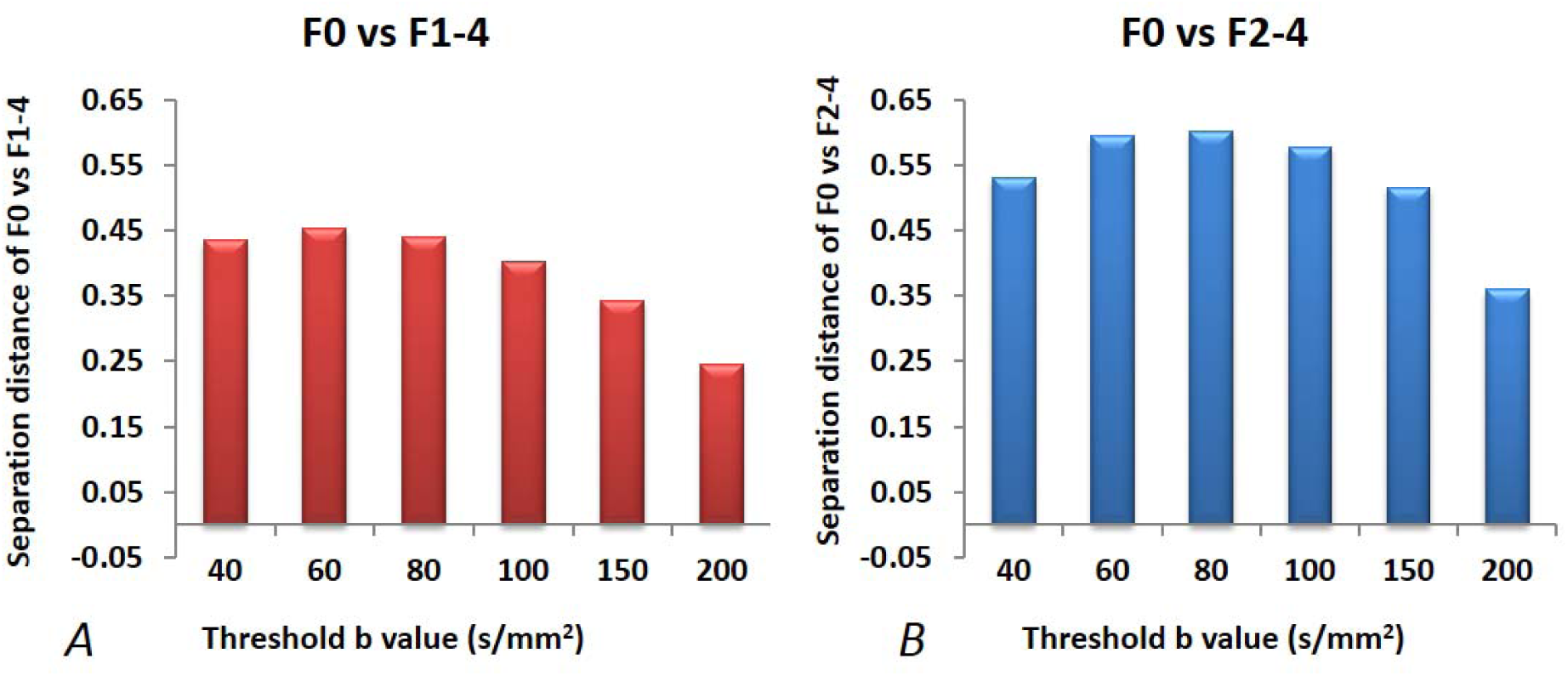
The mean distance (in relative unit) between healthy liver datapoints and fibrotic liver datapoints in 3-dimensional space with threshold *b*-value ranges between 40 -200 s/mm^2^. This distance in x-axis is the sum of the mean distance of F0 liver datapoints to the separation plane and the mean distance of F1-F4 liver datapoints, or F2-F4 liver datapoints, to the separation plane.

**Figure 4.**
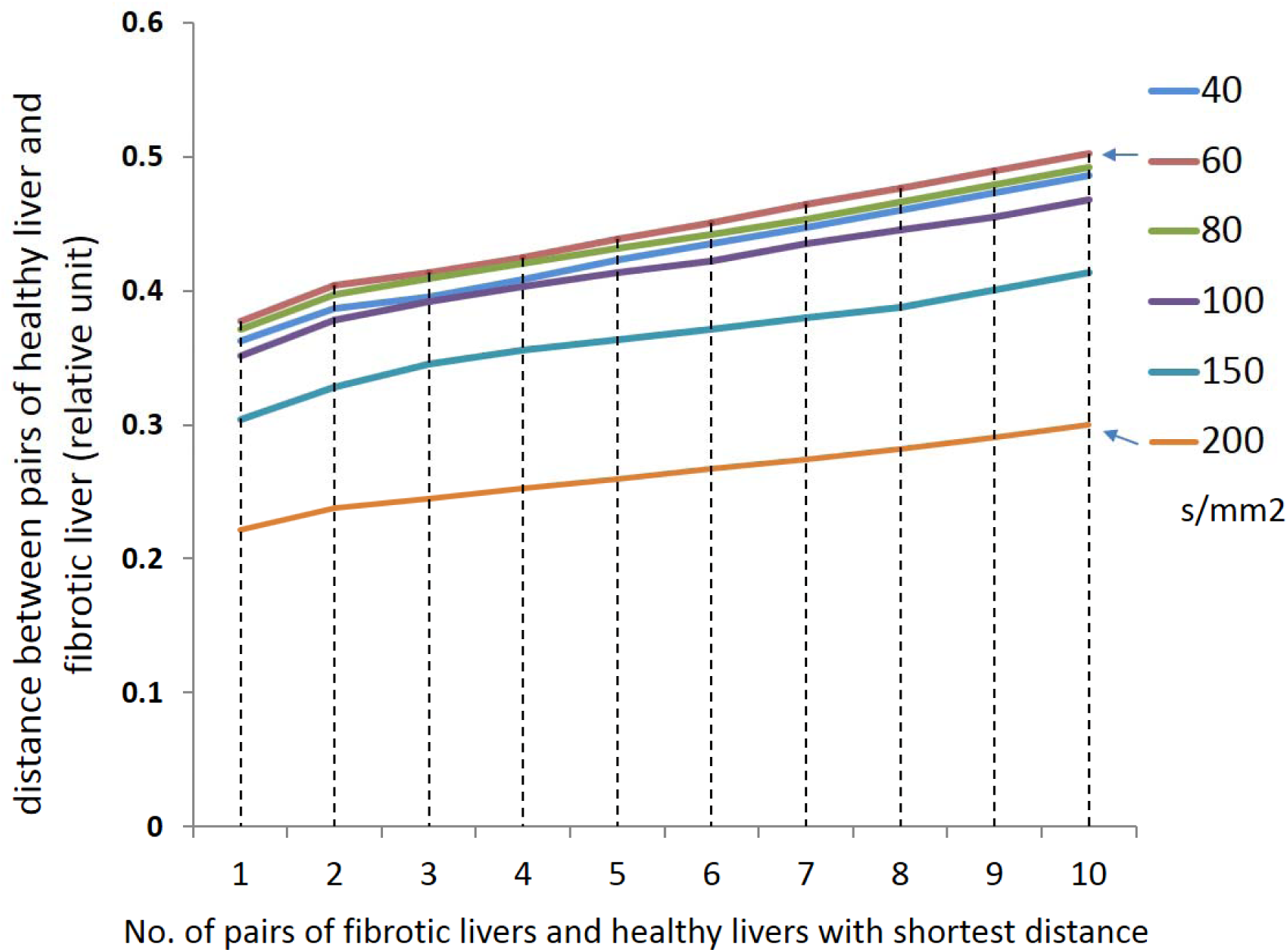
The distance (in relative unit) between healthy liver datapoints and fibrotic liver datapoints in 3-dimensional space with threshold *b*-value ranges between 40 -200 s/mm^2^ (different color lines). The result for first pair of fibrotic liver and healthy liver with shortest distance is denoted in 1; the result for the pair of fibrotic liver and healthy liver with the second shortest distance is denoted in 2; and the result for the pair of fibrotic liver and healthy liver with the tenth shortest distance is denoted in 10. In all cases, *b*-value= 60 s/mm^2^ provides largest mean distance.

**Figure 5.**
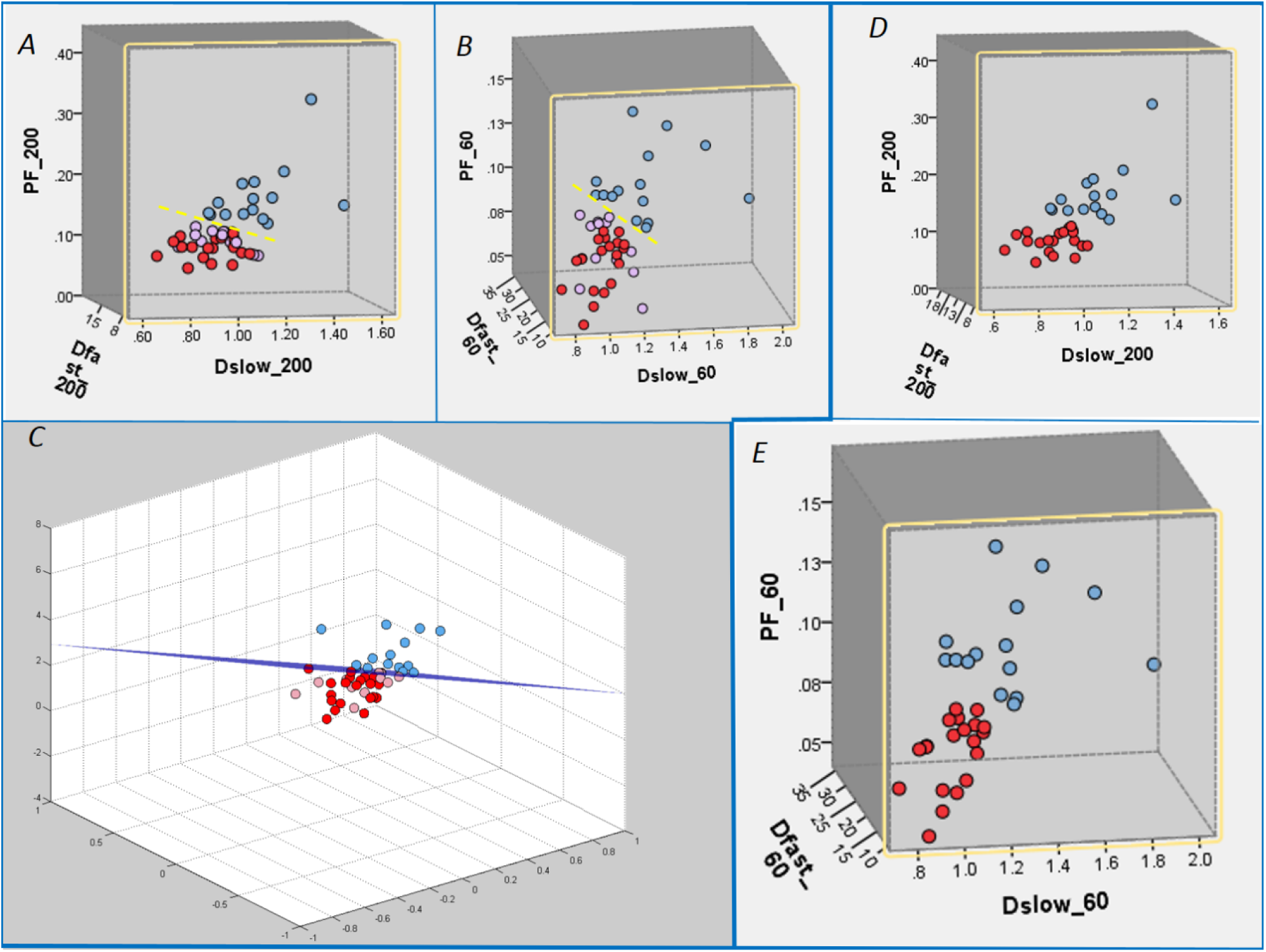
Three-dimensional display of healthy volunteer group (blue balls), F1 patient group (pink balls), and F2–3 patient group (red balls). Each ball represents one participant. The differentiation of the volunteer group and the patient group can be better visualized by rotation in a three-dimensional space (dotted yellow line or purple plane). A: threshold *b*-value=200 s/mm^2^; B: threshold *b*-value=60 s/mm^2^; C: threshold *b*-value=60 s/mm^2^ and the separation plane between healthy livers and fibrotic livers is presented. D &E: threshold *b*-value=200 s/mm^2^ and 60 s/mm^2^ respectively with F1 patients excluded. Threshold *b*-value=60 s/mm^2^ allows better separation between healthy livers and patients.

## Discussion

The recent study of Wáng *et al* reported the possibility that IVIM DWI technique may be able to detect early stage liver fibrosis with good sensitivity and specificity [12]. The current study represents the on-going efforts on optimizing image analysis to further improve the diagnostic performance of liver IVIM DWI. With the typical segmented-unconstrained IVIM algorithm to separate the liver parenchyma perfusion and diffusion compartments, this study empirically demonstrated a higher threshold *b*-value was associated with flatter Dslow curve (i.e. lower Dslow value) and lead to higher PF measurement. On the other hand, a lower threshold *b*-value led to Dslow curve containing more perfusion compartment, and therefore higher Dslow measurement and lower PF measurement (Fig 6). Only including low *b*-values which correlate to the initial sharp signal decay led to higher Dfast measurement (Fig 6). Compared with the previously used threshold *b*-value of 200 s/mm^2^, in this study *b*-value of 60 s/mm^2^ increases the mean distance between healthy liver datapoints and fibrotic liver datapoints in 3-dimensional space by 84.5%. If only considering the separation of significant healthy livers (F0) vs. significant fibrotic livers (F2-4), then the choose of b=value of 80 s/mm^2^ increases mean distance between healthy liver datapoints and fibrotic liver datapoints in 3-dimensional by 65.4%. Classification and regression tree also confirmed improved sensivity and specificty when threshold *b*-value of 60 s/mm^2^ was used as compared with threshold *b*-value = 200 s/mm^2^ (Table 1). This improvement is expected to be important for detecting patients who may have milder liver fibrosis than the F1 cases presented in this study. Additionally, it has been shown previously that for borderline cases, all three IVIM parameters of PF, Dslow and Dfast contribute to the differentiation between healthy livers and fibrotic livers [12].

**Figure 6.**
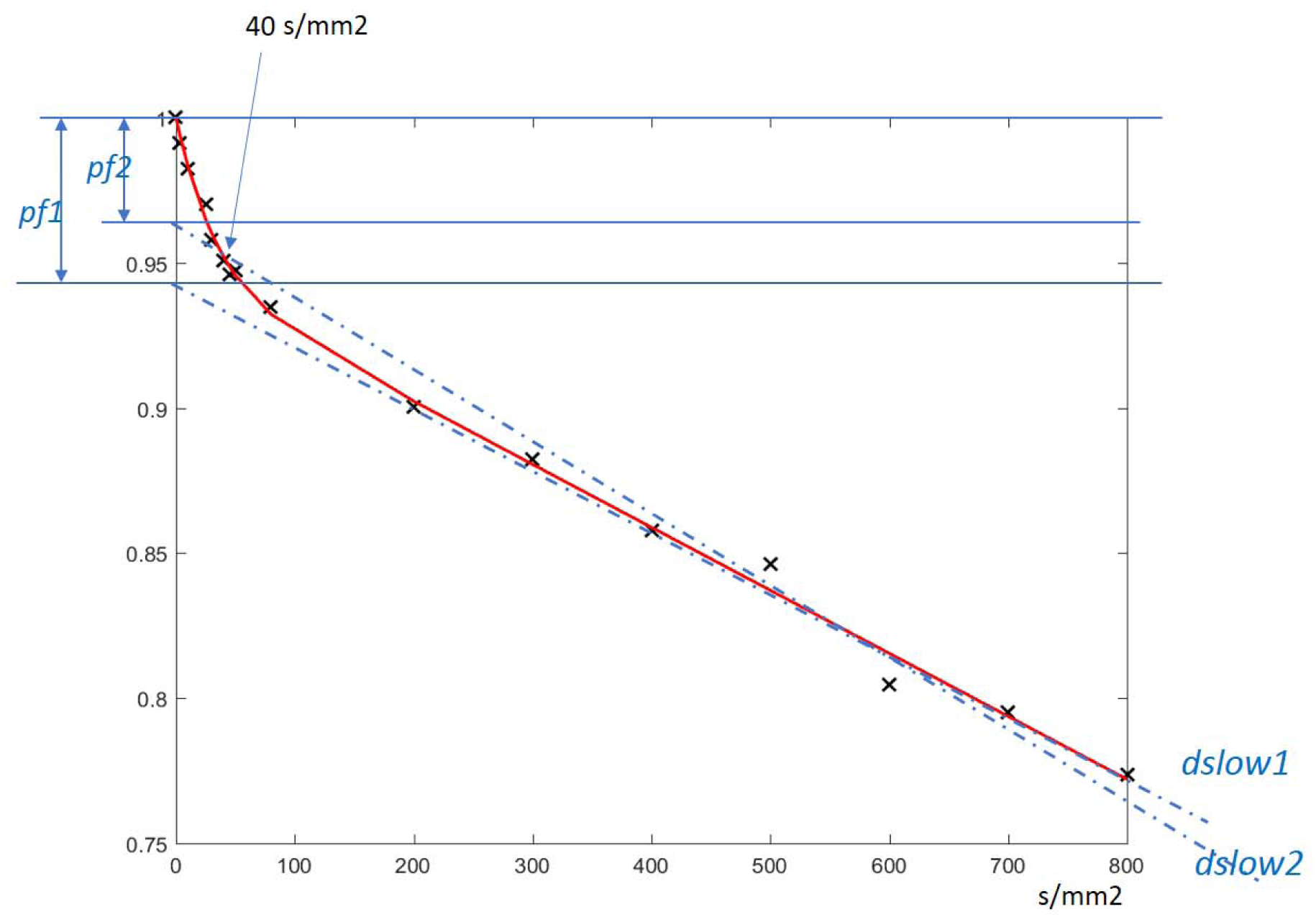
A typical signal intensity vs. *b*-value curve acquired at a 3 Tesla scanner with 16 *b*-value of 0, 3, 10, 25, 30, 40, 45, 50, 80, 200, 300, 400, 500, 600, 700, 800 s/mm^2^ (note more *b*-values were used than the study in this paper). The biexponential decay trend is well presented. It can be seen that when a higher threshold *b*-value (such as *b*=200 s/mm^2^) is used, the Dslow value (slope of *d1* line) will be smaller than when a lower threshold *b*-value (such as *b*=40 s/mm^2^) is selected (slope of *d2* line). On the other hand, with segmented-unconstrained analysis, the computed PF is smaller when a higher threshold *b*-value is used (height of *f1*) than when a lower threshold *b*-value is used (height of *f2*).

Theoretically, there is no ground truth which is the most suitable threshold *b*-value, and we could choose threshold *b*-value=200 s/mm^2^ for PF calculation, and threshold *b*-value=40 s/mm^2^ for Dslow and Dfast calculation. However, this study showed such a selection did not outperform a single *b*-value threshold of 60 s/mm^2^ (Fig 6). Another point of note in this study is that the ROIs were drawn by a trained biomedical engineering graduate student; therefore, the results demonstrated by Wáng *et al* were not dependent on a radiologist’s expertise.

*b*-value=200 s/mm^2^ has so far been a popular choice for IVIM DWI threshold in literature [14-22], as with *b*-value≥200 s/mm^2^ the perfusion compartment contribution to Dslow is considered to be minimal. Recently a number of authors suggested that this threshold *b*-value may be too high as the bi-exponential turning point is generally around 50 s/mm^2^ [7,11]. Other threshold *b*-values have been used in literature, for example, ter Voert *et al* used a threshold of 100 s/mm^2^[24], Dijkstra *et al* used a threshold of 500 s/mm^2^[25]. Our empirical data in this study suggested *b*-value=60 s/mm^2^ should be selected for separating healthy livers and fibrotic livers. This differs from the smallest residuals analysis by Wurnig *et al* [23], where they proposed an optimal threshold *b*-value was 20 s/mm^2^ for liver right robe and 40 s/mm^2^ for liver left robe. However, in Wurnig *et al*’s study only healthy subjects’ data were analyzed. In addition, it is challenging to get reliable measurement of Dfast between *b*-value range of 0-20 s/mm^2^ [30]. In Wáng *et al*’s paper, it was stated that among the IVIM parameters of PF, Dslow and Dfast, PF offers best diagnostic value; and Dslow may suffer from a limited dynamic range and less responsive [12]. The current study demonstrated if a lower threshold *b*-value such as 60 s/mm^2^ is used, then Dslow become more responsive to fibrotic changes (Fig 2).

It is known liver cirrhosis is associated with reduced liver perfusion. Moreno *et al* [9, 10] reported the mean portal flow in healthy subjects was 20.9 mL/min/kg ± 4.1 as opposed to 6.5 mL/min/kg ± 5.6 in patients with cirrhosis. Liver fibrosis is associated with progressive increase in connective tissue, and the increased proportion of collagen fibers is believed to impair Brownian water motion within fibrotic livers. Liver fibrosis may be more associated perfusion compartment (Dfast) reduction than diffusion compartment (Dslow) reduction. The earlier study of Luciani *et al* [15] reported that Dslow do not differ significantly between healthy and cirrhotic livers, and cirrhotic livers is mainly associated Dfast reduction, i.e. reduced perfusion. Luciani *et al* further suggested when considering diffusion weighted imaging in cirrhotic livers, changes in liver architecture may be of less importance than changes in liver perfusion [15]. Recent literature review also showed that among nine patient studies, only Dfast, despite being the least stable, consistently demonstrated liver fibrosis is progressively associated a reduced measurement (Fig 10 of reference 11). Our results showed Dslow value computed from *b*-value range of 40-800 s/mm^2^ is more sensitive to detect fibrotic changes than Dslow value computed from *b*-value range of 200-800 s/mm^2^, therefore Dslow containing more perfusion compartment is more responsive to liver fibrotic changes. This may suggest that study aiming to detect liver fibrosis should include sufficient low *b*-values so to get more reliable perfusion compartment measurement.

The limitations of this dataset have been previously discussed [12]. All causes of chronic liver disease, including viral hepatitis, metabolic, and cholestatic disease, may lead to fibrosis; while all our patients had liver fibrosis due to viral hepatitis-b. Our *b*-value distribution is similar to many previous literature publications, such as Luciani *et al*’s [15], expect that *b*-value=0 was not collected. To include more *b*-values will increase IVIM parameter quantification accuracy [11,23] . While ter Voert *et al* [23] recommended at least 16 *b*-values should be included, Wurnig *et al* [24] and Lemkea *et al* [32] suggested that 10 *b*-values would be an acceptable compromise between accurate measurement, scan time and patient compliance. Without *b*-value=0 computing IVIM parameters with segmented-unconstrained approach, it is expected that both PF and Dfast would have been under-estimated, while Dslow would remain similar. Taking the example of when threshold *b*-value=200 s/mm^2^ is assumed, PF is underestimated by approximately 25% [11, 12]. The echo-planar imaging sequence typically induces bright blood signal, while diffusion weighting induces quick blood signal decay [Fig 7]. This lead to some authors suggested that the IVIM relationship between *b*-value and signal intensity is better fitted with an tri-exponential decay model [32], some authors choose to abandon *b*-value=0 s/mm^2^ for diffusion image series analysis.

**Figure 7.**
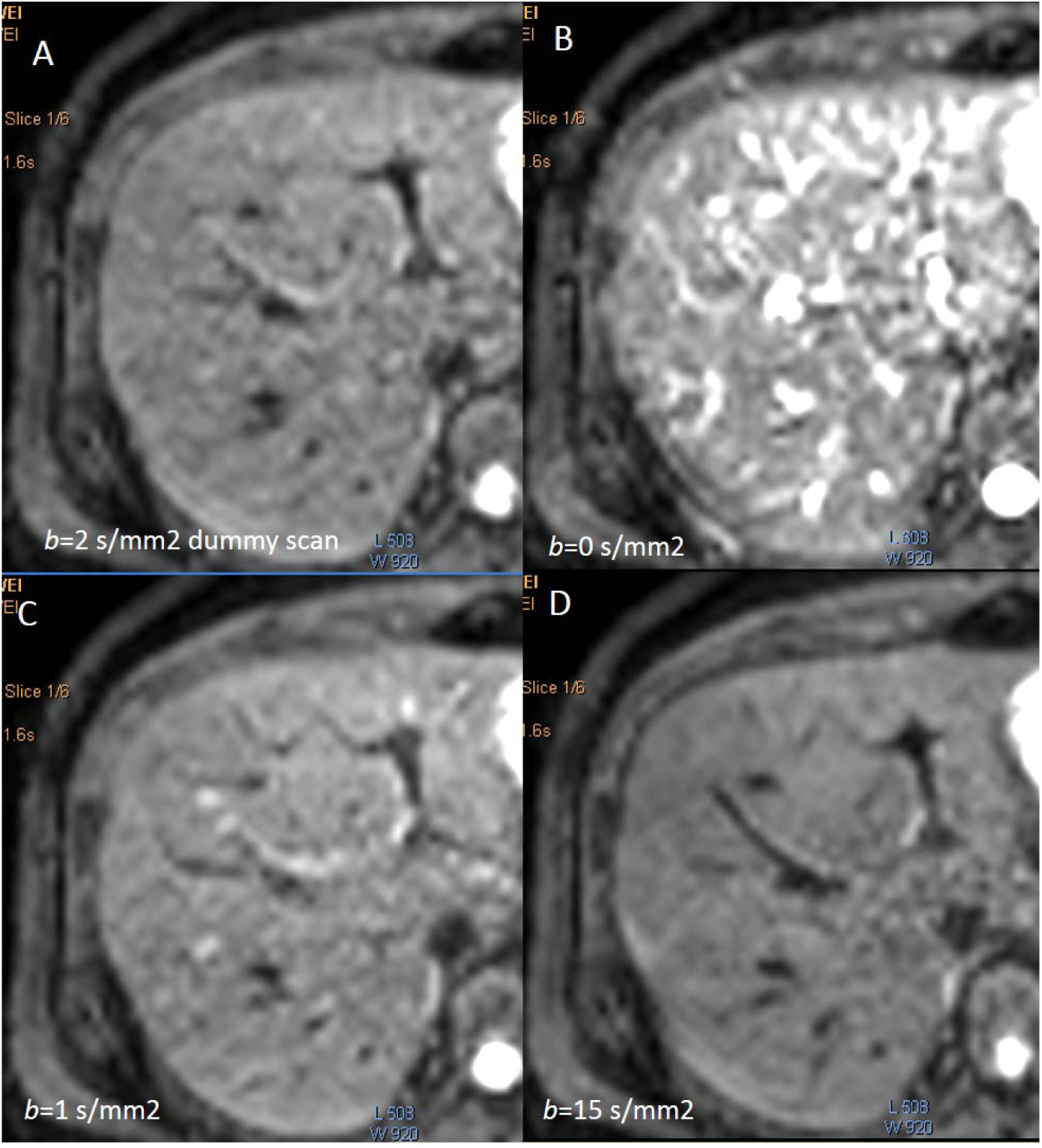
Typical liver EPI (echo planar imaging) diffusion images of *b*-value of 2 (dummy scan), 0, 1, and 15 (s/mm^2^, acquired with the same 1.5 Tesla scanner used for the study in this paper). *b*-value= 0 image (B) is associated with extensive bright blood signal.

There are a number of ways the current data analysis can be further improved. Respiratory motion is major contributor of variation of quantitative liver MR imaging [11, 33], it is expected that motion correction can further improve the reliability of IVIM quantification [34, 35]. Another possibility will be poorly fitted pixels are statistically removed [36]. Finally, IVIM technique can be incorporated to multiparameter diagnosis with the use of Bayesian prediction, incorporating relevant findings from the available methods such as IVIM DWI, liver T1rho, and elastography [37-41].

In conclusion, this study concurs with previous reports that liver fibrosis is more associated perfusion compartment (Dfast) reduction than diffusion compartment (Dslow) reduction. For both healthy livers and fibrotic livers, a higher threshold *b*-value is associated with overall higher PF measurement; while a lower threshold *b*-value leads to overall higher Dslow measurement and higher Dfast measurement. The dependence of PF, Dslow, and Dfast on threshold *b*-value differs between healthy livers and fibrotic livers; with the healthy livers showing a higher degree of dependence. Compared with commonly used threshold *b*-value=200s/mm^2^, threshold *b*-value=60 s/mm^2^ improves differentiation between healthy livers and fibrotic livers.

